## Supplementary Information for "A complex feature-based representation of vocalizations emerges in the superficial layers of primary auditory cortex"

Supplementary material consists of one supplementary figure.

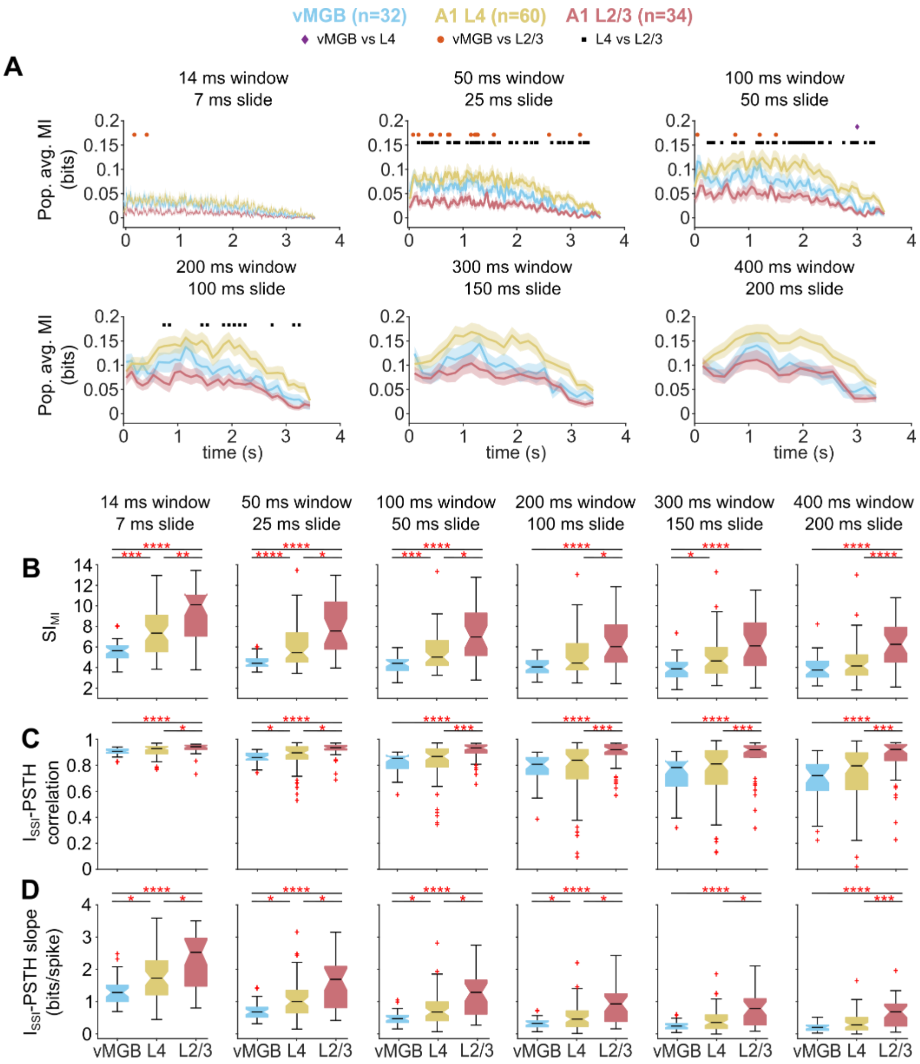


**Supplementary Figure 1: Information analyses at multiple window sizes.**

**(A)** Population average of MI as a function of time in vMGB (blue), A1 L4 (yellow), and A1 L2/3 (red) neurons. Lines correspond to means and shading to 1 s.e.m. Colored dots represent results of statistical testing ($p<0.05$; two-sided t-test with FDR correction for multiple comparisons). Distributions of **(B)** $SI_{MI}$ **(C)** ISSI – PSTH correlation coefficients and **(D)** ISSI – PSTH slopes for vMGB, A1 L4 and A1 L2/3 neurons at all considered window sizes. Asterisks correspond to: *: $p<0.05$, **: $p<0.01$, ***: $p<0.005$, ****: $p<0.001$ (Kruskal-Wallis test with posthoc Dunn-Sidak tests)
